# CIEVaD: a lightweight workflow collection for rapid and on demand deployment of end-to-end testing of genomic variant detection

**DOI:** 10.1101/2024.06.21.600013

**Authors:** Thomas Krannich, Dimitri Ternovoj, Sofia Paraskevopoulou, Stephan Fuchs

## Abstract

The identification of genomic variants has become a routine task in the thriving age of genome sequencing. Particularly small genomic variants of single or few nucleotides are routinely investigated for their impact on an organism’s phenotype. Hence, precise and robust detection of the variants’ exact genomic location and change in nucleotide composition is vital in many biological applications. Although a plethora of methods exist for the many key steps of variant detection, thoroughly testing the detection process and evaluating its results is still a cumbersome procedure. In this work, we present a collection of trivial to apply and highly modifiable workflows to facilitate the generation of synthetic test data as well as to evaluate the accordance of a user-provided set of variants with the test data.

**Availability:** The workflows are implemented in Nextflow and are freely available and open-source at https://github.com/rki-mf1/cievad under the GPL-3.0 license.

## Introduction

Since the dawn of DNA sequencing technologies and genomic diagnostics, more and more light has been shed on the evolution and variability of genomes. Over the past decades, changes in the nucleotide composition and structure have been observed and extensively studied in a variety of organisms. A common observation when comparing genomes over time, across individuals or in different cell types are alterations of single or multiple nucleotides. In many studies, observations of these alterations, termed genomic variants, have been associated with diseases, evolutionary processes and biodiversity (1, 2).

The corollary and benefit of a comprehensive understanding of genomic variants has driven immense efforts to collect and analyze genomic data. A plethora of bioinformatics software has been released to accurately quantify and describe the characteristics of genomic variants in a single or multiple genomes (3). For such a software product to prevail over time, it is often subject to long-term update cycles, compared to novel competing software in the field, and analyzed for technical errors by the community. These challenges of continuous software development require robust testing and evaluation of results (4). In the context of genomic variant detection, a bioinformatics software needs to perform well under a variety of technical setups and the set of reported genomic variants should comply with a known ground truth.

Several existing applications provide functionalities to perform individual steps towards end-to-end testing of genomic variant detection. Mason (5) and ART (6) are applications to generate synthetic sequencing data of different sequencing technologies. PICARD (7) and BCFtools (8) are established tool suites for manipulating high-throughput sequencing data and variant call format (VCF) files. Krusche et al. (9) proposed a framework describing best practices for benchmarking small germline variants, which led to the highly recognized precisionFDA challenge (10). The same author released the widely community-accepted *hap*.*py* tools for variant set comparison. This tool suite implements elaborate methods to address complex cases where representational differences between sets of variants cannot be trivially fixed. Another commonly used alternative of *hap*.*py* is RTG Tools (https://github.com/RealTimeGenomics/rtg-tools). Vcfdist (11) is another evaluation tool with a focus on using local phasing information. The recent NCbench (12) is an elaborate benchmark platform for sets of small genomic variants. It has a competitive advantage in interactivity and visualization. However, none of these individual applications orchestrates an all-in-one solution for flexible, on demand and easy-to-install end-to-end testing of genomic variant detection. To this end, we introduce *CIEVaD* (abbreviation for <u>C</u>ontinuous <u>I</u>ntegration and <u>E</u>valuation of <u>Va</u>riant <u>D</u>etection), a lightweight workflow collection designed for rapid generation of synthetic test data and validation of software for genomic variant detection. CIEVaD is implemented in Nextflow (13), a workflow management framework for streamlined and reproducible data processing. Using Nextflow additionally avoids manual installation efforts for the user via the deployment of software packages.

## Methods

The CIEVaD workflow collection contains two main workflows. The haplotype workflow generates synthetic data whereas the evaluation workflow examines the accordance between sets of genomic variants. In the following, the term *synthetic* is used synonymously for *in-silico generated*.

### Haplotype workflow

The haplotype workflow provides a framework for data generation. Synthetic data can be used for genomic variant calling, and the data generation scales computationally to many individuals. Also, different types of sequencing data can be specified. To get started with the data generation the one input parameter strictly required by the user is a reference genome. With only a given reference genome, the haplotype workflow generates a new haplotype sequence of the entire reference genome, a set of synthetic genomic variants (henceforth referred to as *truthset*), and synthetic genomic reads. The truthset comprises single-nucleotide variants (SNVs) and short insertions and deletions (indels) of at most 20 nucleotides. The maximum length of indels as well as the frequency of both variant types can be adjusted via workflow parameters. Since the variants of the truthset are homozygous, the alternative-allele ratio for each variant is defined by 1− *ϵ*, where *ϵ* is drawn from the read-specific error distribution.

In the initial step, the workflow indexes the given reference genome. Next, CIEVaD uses the Mason variator (5) for a given number *n* of individuals (default is *n* = 3). The result of this step is a haplotype sequence and truthset per individual. Here, a haplotype sequence is a copy of the reference genome with small genomic variants of the truthset inserted at pseudo-random locations. In other terms, the reference genome and a haplotype sequence differ by the variants of the corresponding truthset. The final step is the generation of synthetic genomic reads. Depending on the specified type of sequencing data (default is *NGS*) either the Mason simulator generates pairs of genomic reads or PBSIM3 (14) generates a set of synthetic long-reads. In both cases, the workflow returns an alignment of the reads to the reference genome, e.g., for a manual inspection of the genomic variants and sequencing artifacts in the reads. Note that the data of each individual is computed in asynchronous parallel Nextflow processes which scale effortlessly with additional CPU threads or compute nodes.

### Evaluation workflow

The objective of the evaluation workflow is to assess how successfully a third-party tool or workflow detects genomic variants. To assess the detection, CIEVaD’s evaluation workflow compares the set of genomic variants generated by the third-party tool or workflow (such a set is further referred to as *callset*) with the corresponding truthset. The only strictly required input of the evaluation workflow is a set of callsets in the Variant Call Format (VCF), either given as a folder path or a sample sheet. The output are reports about the accordance between corresponding truth- and callsets.

The evaluation workflow consists of only two consecutive steps. First, the open-source tool suite *hap*.*py* by Illumina Inc is used to compare all truthsets with their corresponding callsets. In fact, its included submodule *som*.*py* identifies the number of correctly detected (TP), missed (FN) and erroneously detected (FP) variants. By using *som*.*py* this comparison of the variants neglects the genotype information and only checks their position in the genome as well as the nucleotide composition. This default behaviour is chosen since CIEVaD was initially developed for haploid pathogens where the presence of a variant itself reflects the genotype. Using the TP, FP and FN counts, *som*.*py* reports the precision, recall and F1-score of the variant detection process that yields the callset. The second step of the evaluation workflow is a computation of average scores across the statistics of all individuals.

## Results

To demonstrate the utility of CIEVaD we implemented end-to-end tests (Supplementary material A) for different variant detection software as part of their continuous integration frameworks.

### Assessing variant detection from NGS data as part of SARS-CoV-2 genome reconstruction

We deployed CIEVaD to benchmark the variant detection routine within CoVpipe2 (15). CoVpipe2 is a computational workflow for the reconstruction of SARS-CoV-2 genomes. One sub-process of CoVpipe2 applies the FreeBayes (16) variant detection method to a set of reference-aligned NGS reads.

We implemented a test as part of CoVpipe2’s Github Actions^1^ with the objective to obtain F1-scores for the detection of SNVs and indels. The test consists of seven principal steps:

1. Install Conda and Nextflow
2. Download a reference genome
3. Run CIEVaD hap.nf
4. Run CoVpipe2
5. Prepare input for CIEVaD eval.nf
6. Run CIEVaD eval.nf
7. Check results

In brief, the test runs the CIEVaD haplotype workflow with default parameters, CoVpipe2 with the generated synthetic NGS data, the CIEVaD evaluation workflow with the filtered callsets from CoVpipe2, and finally checks whether the F1-scores of the SNV and indel detection decreased compared to previous test scores. This rapid deployment of CIEVaD (v0.4.1) benchmarks the variant callsets of CoVpipe2 (v0.5.2) with F1-scores of 0.97 and 0.91 for SNVs and indels, respectively. The full table of results of this evaluation is in Supplementary material B.

### Assessing the variant detection from long-read data as part of nanopore sequencing data analysis

In order to deploy CIEVaD for a different than the previous genomic data type we generated long-reads with hap.nf using the *–read_type* parameter (see Supplementary material C). The read type parameter invokes additional default parameters of the haplotype workflow that are tailored to a high-coverage long-read WGS experiment of the SARS-CoV-2 genome. With this setup, the haplotype workflow and its internal long-read module (14) generates a dataset with an average of 500-fold read coverage, a per base accuracy of approximately 90% (Supplementary material D) and an error distribution model trained on genomic data from Oxford Nanopore Technologies.

We used the synthetic long-read dataset and their ground truth variants to test the variant detection of the poreCov (17) data analysis pipeline. Supplementary material C shows how we adjusted poreCov (v1.9.4) to process the synthetic long-read dataset and, subsequently, how the evaluation workflow of CIEVaD verifies poreCov’s results. Our evaluation of poreCov (Supplementary material E) shows F1-scores of 0.95 and 0.73 for SNVs and indels, respectively.

## Conclusions

We introduce CIEVaD, an easy-to-apply tool suite to assess small variant detection from short and long-read datasets. CIEVaD is modular, extensible, scaleable, requires no manual installation of internal software and operates entirely on standard bioinformatics file formats. We showed that the workflows of CIEVaD enable a rapid deployment of end-to-end tests including generation of synthetic genomic data and evaluation of results from third-party variant detection software.

With the workflow design, open-source policy, file formats, software packaging, and documentation, we aim to comply with the PHA4GE best practices for public health pipelines (https://github.com/pha4ge/public-health-pipeline-best-practices/blob/main/docs/pipeline-best-practices.md). Within the scope of this work, we did not implement advanced continuous integration strategies for third-party software, but it is up to the user to apply arithmetic operations or use test frameworks around the results of CIEVaD. It should be mentioned here that CIEVaD is not restricted to synthetic data, the evaluation workflow also works with curated variant callsets from real sequencing data. Current limitations of CIEVaD comprise the zygosity and type of the variants. As the project and workflows were initially intended for viral pathogens, we have not implemented an option to generate heterozygous variants. For the same reason, we have not provided an option to generate synthetic structural variants. Also, structural variants require more sophisticated algorithms for variant set comparison. Features for heterozygous and structural variants remain subject to the users’ demand.

## ACKNOWLEDGEMENTS

The authors would like to thank Namuun Battur for testing early versions of the workflow collection and Marie Lataretu for her feedback on using CoVpipe2 and proofreading the manuscript.

## AUTHOR CONTRIBUTIONS

TK implemented CIEVaD and the tests of third-party software. DT contributed implementations for additional features in CIEVaD. SP and SF supervised the project. TK wrote the manuscript. All authors read and approved the manuscript.

## Footnotes

1 https://github.com/rki-mf1/CoVpipe2/actions/workflows/VariantCalling.yml

