## Supplemental material for "CIEVaD: a lightweight workflow collection for rapid and on demand deployment of end-to-end testing of genomic variant detection"

### Supplementary material

#### A. Workflow scheme of an end-to-end test using CIEVaD.

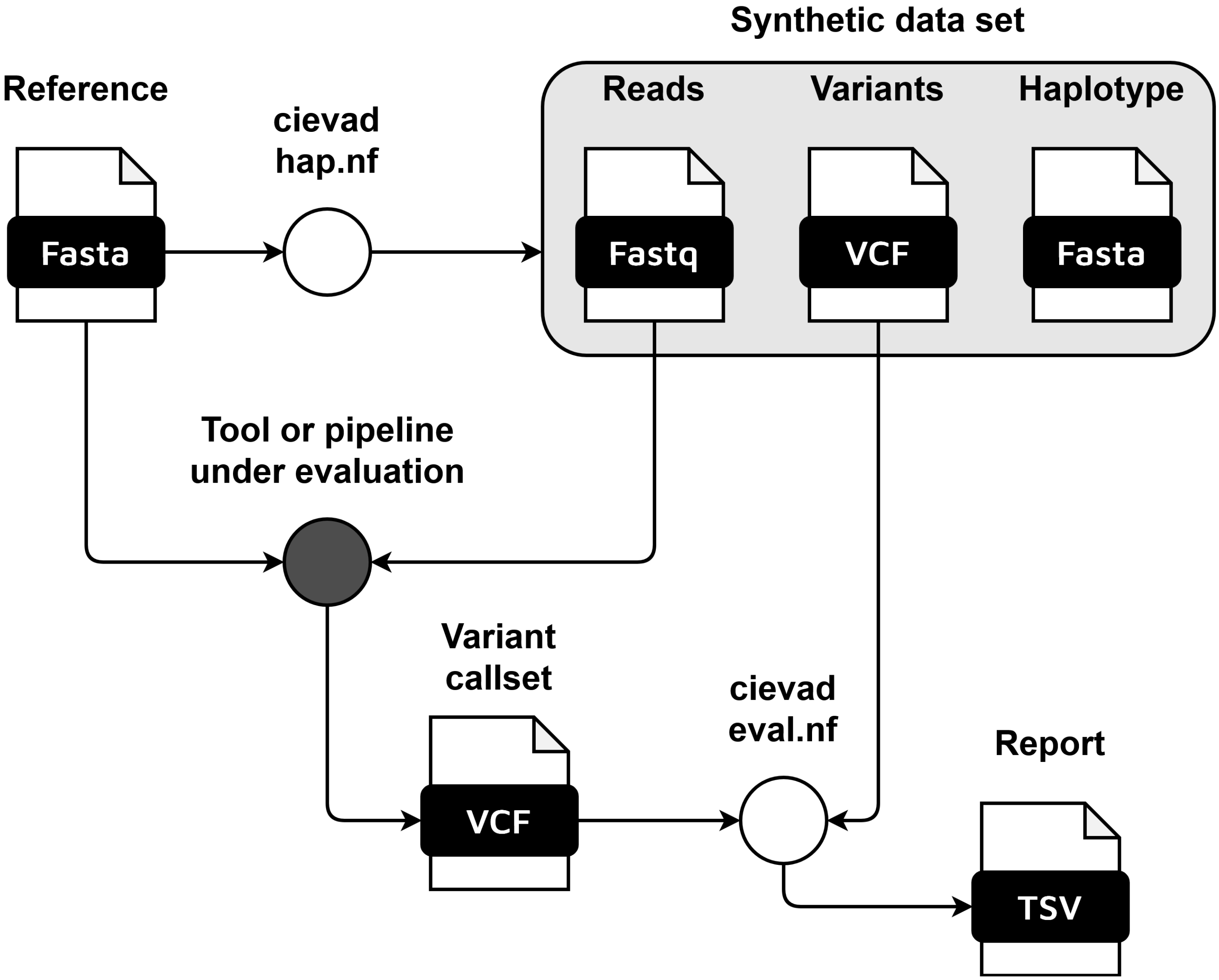


**Fig. 1.** General overview of an end-to-end test using CIEVaD. The document pictograms show the data and file formats that are passed throughout the testing process. CIEVaD can be deployed around any variant detection software or pipeline as long as it returns a variant callset in Variant Call Format (VCF).

**B. Pivotal results from the evaluation of CoVpipe2.** The table shows the precision and recall statistics of the variant detection process of CoVpipe2 (v0.5.2) evaluated with CIEVaD (v0.4.1).

| Type | Total in truthset | Total in query | TP | FP | FN | Recall | Precision | F1-score |
|---|---|---|---|---|---|---|---|---|
| indels | 158.67 | 134.33 | 133.67 | 0.67 | 25 | 0.84 | 0.996 | 0.91 |
| SNVs | 287.33 | 268.67 | 268.67 | 0 | 18.67 | 0.93 | 1 | 0.97 |

**Table 1.** Results of the CIEVaD evaluation workflow using CoVpipe2's variant callsets. All values are average values across three synthetic individuals. The total in truthset, total in query, TP, FP and FN are the averages of the individuals' case counts, hence from $\mathbb{R}_0^+$. Precision, recall and F1-score are floating point numbers from the interval $[0, 1]$. Here, the term *query* is synonymous for callset, as used by the hap.py tools suite. The F1-score is the harmonic mean of precision and recall.

**C. Shell commands used to test poreCov using CIEVaD.** These shell commands were executed on a Linux compute node with an Ubuntu 20.04.6 LTS operating system and equipped with an AMD EPYC 9534 64-Core Processor. The working environment has the *conda* package management system and *singularity* container virtualization software pre-installed.

```
#!/bin/bash
#run synthetic data generation
conda create -n nextflow -c bioconda nextflow=23.10.1
conda activate nextflow
git clone https://github.com/rki-mf1/cievad.git && cd cievad
nextflow run hap.nf -profile local,conda --read_type ont
conda deactivate

#run poreCov
conda create -n nextflow-2104 -c bioconda nextflow=21.04.0
conda activate nextflow-2104
nextflow run replikation/poreCov -r 1.9.4 -dsl2 \
    --fastq "results/simulated_hap*.fastq" \
    -profile local,singularity --cores 2 --max_cores 8
conda deactivate

#run callset evaluation
echo "index,truthset,callset" > results/sample_sheet.csv
for i in {1..3}
do
    echo "${i},${PWD}/results/simulated_hap${i}.vcf,\
    ${PWD}/results/3.Lineages_Clades_Mutations/simulated_hap${i}\
    /SNP_simulated_hap${i}.pass.vcf" >> results/sample_sheet.csv
done
conda activate nextflow
nextflow run eval.nf -profile local,conda --sample_sheet results/sample_sheet.csv
column -s ',' -t results/summary.sompy.stats.csv | less -S
```

##### D. Synthetic long-read data from the CIEVaD haplotype workflow.

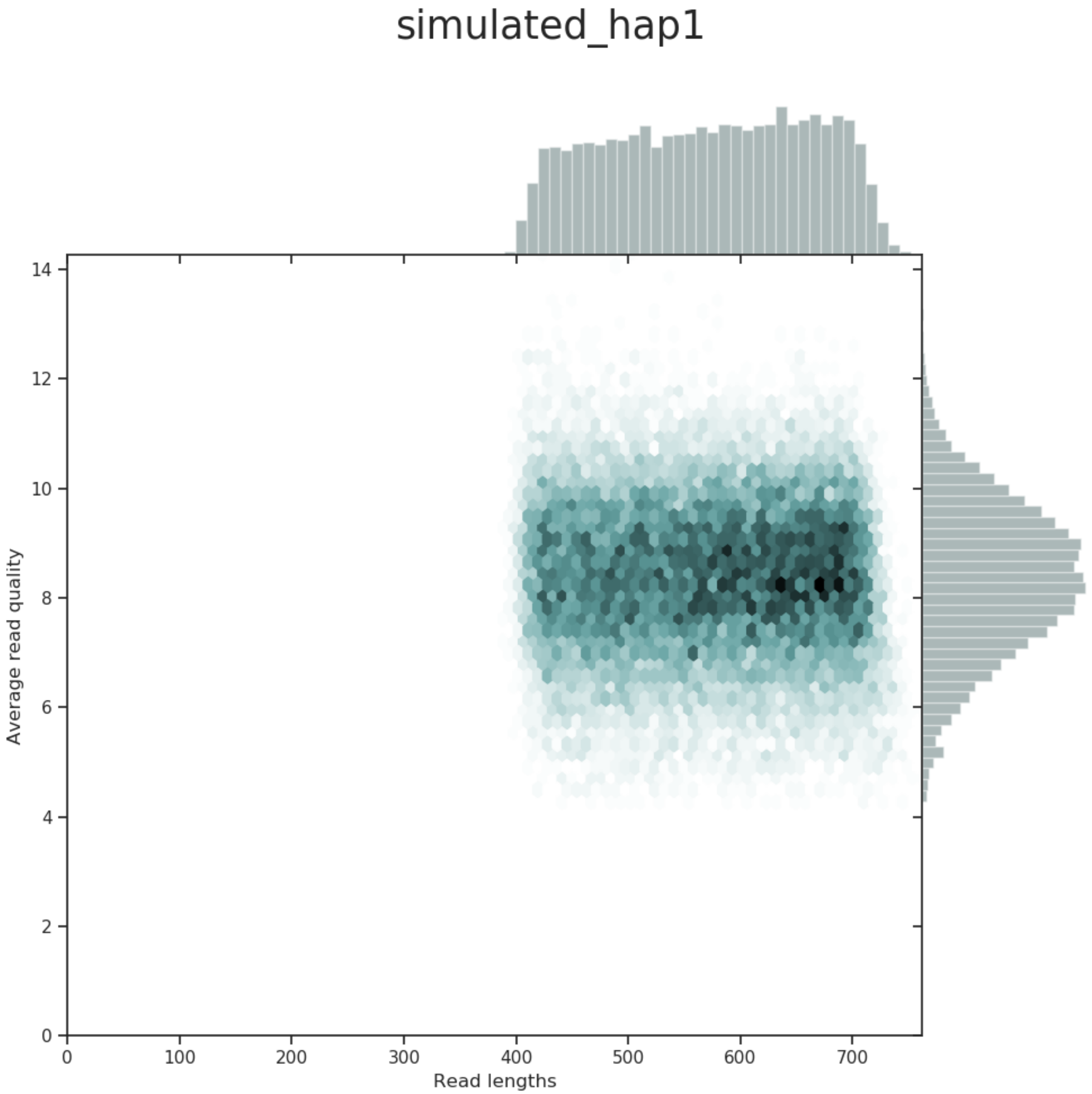


**Fig. 2.** Heat-map of read lengths versus average read quality. The visualization is automatically generated with Nanoplot as part of the poreCov workflow. The long reads were generated with PBSIM3. The length interval of the generated reads is chosen according to the read filter criteria of poreCov. The average read quality provides a challenging yet manageable signal-to-noise ratio for testing.

**E. Pivotal results from the evaluation of poreCov.** The table shows the precision and recall statistics of the variant detection process of poreCov (v1.9.4) evaluated with CIEVaD (v0.4.1).

| Type | Total in truthset | Total in query | TP | FP | FN | Recall | Precision | F1 |
|---|---|---|---|---|---|---|---|---|
| indels | 158.67 | 166 | 119 | 47 | 39.67 | 0.75 | 0.72 | 0.73 |
| SNVs | 287.33 | 263.33 | 261 | 2.33 | 26.33 | 0.91 | 0.99 | 0.95 |

**Table 2.** Results of the CIEVaD evaluation workflow using poreCov's variant callsets. All values are average values across three synthetic individuals. The total in truthset, total in query, TP, FP and FN are the averages of the individuals' case counts, hence from $\mathbb{R}_0^+$. Precision, recall and F1-score are floating point numbers from the interval $[0, 1]$. Here, the term *query* is synonymous for callset, as used by the hap.py tools suite. The F1-score is the harmonic mean of precision and recall.
